# Developmental time course of SNAP-25 isoform regulation of hippocampal long-term synaptic plasticity and hippocampus-dependent learning

**DOI:** 10.1101/542928

**Authors:** Katisha R. Gopaul, Muhammad Irfan, Omid Miry, Linnea R. Vose, Alexander A. Moghadam, Tomas Hökfelt, Christina Bark, Patric K. Stanton

## Abstract

SNAP-25 is essential in activity-dependent vesicle fusion and neurotransmitter release in the nervous system. During development and adulthood, SNAP-25 appears to have differential influences on long- and short-term synaptic plasticity in the hippocampus. The involvement of SNAP-25 in this process may be altered by two different splice variants expressed in adolescences versus adulthood in hippocampal neurons. This study suggests that the adolescent isoform, SNAP-25a can contribute to developmental regulation of the expression of LTD and LTP. In mice deficient in SNAP-25b, the adult isoform, Schaffer collateral-CA1 synapses showed slower release kinetics, reduced initial release probabilities, decreased LTP and enhanced LTD at 1 month. By 4 months of age, when mice have fully developed in the absence of SNAP-25b, the gene targeted mice appear to have compensated for the lack of the adult SNAP-25b isoform. Moreover, hippocampal-dependent training reversed reductions in LTP, but not LTD, seen at 1 month. In 4 month old adult mice, training prevented the compensatory reversal of LTD that had been observed prior to training. These findings support the hypothesis that immature SNAP-25a plays a strong role in the expression of plasticity at Schaffer collateral-CA1 synapses in adolescent mice, but compensatory mechanisms that reverse alterations in synaptic plasticity once mice reach adulthood.

## Introduction

Synaptosomal Associated Protein-25 (SNAP-25) is one of 3 core **SNA**P (Soluble N-ethylmaleimide–Sensitive Factor Attachment Protein) **Re**ceptor (**SNARE**) proteins that participate in the fusion of membrane vesicles with the plasma membrane. SNARE complex proteins interact to overcome the energy barrier for plasma membrane fusion[1–3]. Protein-protein interactions between these SNARE fusion proteins and additional proteins can modify the binding properties and alter vesicle fusion[2,3]. While the other core SNARE proteins, synaptobrevin and syntaxin, are embedded in their respective membranes, SNAP-25 tethers to the postsynaptic membrane via palmitoylation, a dynamic acylation, and binds in the SNARE complex by forming a coiled coil quaternary structure where alpha helices of all three proteins wrap around each other. In the nervous system, SNAP-25 plays a major role in neurotransmitter release, and during development, SNAP-25 has also been suggested to play a role in promoting neurite outgrowth[4,5].

Alternative splicing is a key mechanism for increasing the diversity profile of most protein coding genes. More than 95% of multi-exon genes in higher eukaryotes go through alternative splicing[6,7], and SNAP-25 utilizes this process to produce two variants, SNAP-25a, which is the predominant isoform in neurons of adolescent mice, and SNAP-25b, which becomes the dominant form as synapses mature into adulthood[8]. The two splice variants differ by only 9 amino acids[8,9]. However, SNAP-25a still is the major isoform in endocrine, neuroendocrine and selected neuronal populations throughout life. Moreover, the differences between SNAP-25a and SNAP-25b lie within the cysteine-rich linker region, in the middle of the protein, spacing the two amphiphatic helices participating in SNARE complex formation and within the different exon 5’s[5]. However, the major difference in release efficiency of these isoforms are not purely related to the four cysteine residues that tether the palmitoyl side chains that anchor the protein to the cytosolic face of presynaptic terminal membranes, but instead, it appears that it can mediate SNAP-25 binding partners at amino acids closer to the C-terminal end[9]. Likely, the different SNAP-25 isoforms change the tertiary structure of the core SNARE complex, and thereby modify the affinities of SNARE-binding partner proteins. Additionally, SNAP-25 isoforms differentially interact with accessory proteins such as Munc 18-1 and Gβγ subtypes[10]. The gene transcript of SNAP-25 contains two tandem repeats of exon 5, 5a and 5b[4]. During mRNA formation, the splicing machinery either splices in exon 5a or exon 5b, corresponding to SNAP-25a or SNAP-25b, respectively. While the phenotypic outcomes of alternate splicing in SNAP-25 are still being elucidated, there are measurable differences in mice expressing only SNAP-25a versus predominantly SNAP-25b[11,12].

Within the first week of life, in the mouse hippocampal formation, SNAP-25a is the dominant form that participates in vesicle fusion and development in the mouse hippocampal formation[7,9,10]. As mice mature, SNAP-25b and overall SNAP-25 levels increase dramatically in the entire brain. By 3 weeks of age, the level of SNAP-25b is more than 50% greater than SNAP-25a. Additionally, between weeks 3-8, during which time synapses complete their maturation, there is a dramatic increase in the total amount of SNAP-25, with SNAP-25a only accounting for about 5 percent of total SNAP-25 mRNA in mouse brain[7].

Interestingly, during the first two weeks of development, rodents exhibit predominantly, if not entirely, metabotropic glutamate receptor (mGluR)-dependent forms of long-term depression of synaptic transmission (LTD), while long-term potentiation (LTP) has yet to appear[13,14]. The timing of this change in relative levels of SNAP-25a to SNAP-25b, mirrors the development up-regulation of N-methyl-D-aspartate receptors (NMDARs), suggesting that SNAP-25a may play a greater role in the expression of mGluR-dependent LTD, while SNAP-25b may be more important for expression of NMDAR-dependent LTD and LTP. It is possible, but still undetermined, whether the switch in expression from SNAP-25a to SNAP-25b confers this ability to express LTD and LTP in older animals.

In young/adolescent mice, the SNAP-25a isoform may facilitate the expression of LTD at Schaffer collateral-CA1 synapses. It is unknown how the amino acid differences between SNAP-25 isoforms produce the functional differences of the SNAP-25a and SNAP-25b protein. However, identifying differential effects of these isoforms on long-term plasticity, during development, is a key point of interest in elucidating their regulation of the balance between LTP and LTD and how it shifts during development.

We recently demonstrated that the developmental switch from SNAP-25a to SNAP-25b in mouse hippocampus differs in timing between males and females[15]. Furthermore, at 4 weeks of age, SNAP-25b-deficient mice demonstrate a significantly reduced LTP and less ability to discriminate between intensities of presynaptic stimuli[15]. Thus, it appears that in hippocampal neurons, the developmental switch to the SNAP-25b isoform is required to cognitive performances related to learning and memory. To further evaluate the role of SNAP-25 on synaptic plasticity and development, we studied SNAP-25b-deficient mice that were genetically modified to only express the adolescent SNAP-25a isoform throughout life, being unable to switch to expressing the adult SNAP-25b form[16]. The developmental time when SNAP-25b displays its most significant increase coincides with a shift from expression of LTD to expression of both LTD and LTP[7,13,14]. Since the switch between isoforms happens between postnatal weeks 1-3, we compared hippocampal slices from one-month old and 4 month old mice, to assess the early developmental and later adult impact of an absence of SNAP-25b on long-term synaptic plasticity and learning. Improving our knowledge of presynaptic mechanisms of plasticity, the molecules that mediate these changes and the outcomes they have on learning and memory can help identify therapeutic targets for disorders associated with synaptic plasticity dysfunction.

## Methods

### Hippocampal Slice Preparation and Recording

Coronal brain slices were used to analyze hippocampal Schaffer collateral-CA1 synapses. Brains were removed from male and female mice decapitated under deep isoflurane anesthesia, and immersed in ice-cold (2-4°C) oxygenated sucrose-containing artificial cerebrospinal fluid (ACSF) in mM: NaCl 87, NaHCO_3_ 25, NaH_2_PO_4_ 1.25, KCl 2.5, CaCl_2_ 0.5, MgCl_2_ 7, D-glucose 25, Sucrose 75, (saturated with 95% O2/5% CO2, pH 7.4). The cerebellum and frontal lobes were removed, and brains hemisected and each half mounted on a metal stage with cyanoacrylate glue and again submerged in ice-cold sucrose-containing ACSF. 350-400μm thick coronal slices were cut with a vibratome (Leica model VT1200S), placed in an interface holding chamber at 32°±2°C for at least 30 minutes, and then transferred to room-temperature ACSF in mM: NaCl 126, NaHCO_3_ 26, NaH_2_PO_4_ 1.25, KCl 3, CaCl_2_ 2.5, MgCl_2_ 2, D-glucose 10 (constantly bubbled with 95% O2/5% CO2, pH 7.4) for an additional 30 minutes before start of recording. Slices were maintained at room temperature until being moved to an interface recording chamber and maintained thereafter at 32°±2°C for the remainder of the experiment.

Slices were continuously perfused at a rate of 2 ml/min with oxygenated normal ACSF (Gibson Minipuls 3). Postsynaptic excitatory field potentials (fEPSPs) were evoked in field CA1 by activation of Schaffer collateral axons using a stainless steel bipolar stimulating electrode placed in the stratum radiatum of CA3 pyramidal neuron axons. A thin-walled glass microelectrode was pulled (1-2 megaohm; Flaming/Brown micropipette puller, Model P-97, Sutter Instrument) and filled with normal ACSF. The borosilicate micropipette was position in CA1 apical dendritic arbors to record fEPSPs. A constant current stimulation was applied by an ISO-Flex isolator driven by a Master-8 programmable pulse generator (A.M.P.I, Jerusalem, Israel) and recorded at half maximal fEPSP amplitude once every 30 seconds. Recordings were amplified by Multiclamp 700B Axon Instruments (Molecular Devises), acquired by a National Semiconductor AD board, and analyzed using SciWorks software (DataWave Technologies) by calculating the maximum slope within 20-80% of the maximum fEPSP initial negative slope. At least 15-minute stable baselines were recorded at 0.033Hz prior to application of either high frequency theta burst stimulation (TBS; 4 trains of 10 bursts of 5 pulses each at 100Hz with a 200msec interburst interval, applied at 3 min intervals) to induce LTP, or a single train of low frequency stimuli (LFS; 2Hz/10min) to induce LTD. fEPSP slopes were analyzed by Student’s *t*-test, with significance preset to p<0.05. All data points are mean ± SEM.

### Two-photon laser scanning microscopy of FM1-43 vesicular release from Schaffer collateral presynaptic terminals

Fluorescence was visualized using a customized two-photon laser-scanning Olympus BX61WI microscope with a 60x/0.90W water immersion infrared objective lens and an Olympus multispectral confocal laser scan unit. The light source was a Mai-Tai™ laser (Solid-State Laser Co., Mountain View, CA), tuned to 860 nm for exciting Magnesium Green and 820 nm for exciting FM1-43. Epifluorescence was detected with photomultiplier tubes of the confocal laser scan head with pinhole maximally opened and emission spectral window optimized for signal over background. In the transfluorescent pathway, a 565 nm dichroic mirror was used to separate green and red fluorescence to eliminate transmitted or reflected excitation light (Chroma Technology, Rockingham, VT). After confirming the presence of Schaffer collateral-evoked fEPSPs >1 mV in amplitude in CA1 *stratum radiatum*, and inducing LTP, 10 μM 6-cyano-7-nitroquinoxaline-2,3-dione (CNQX) was bath-applied throughout the rest of the experiment to prevent synaptically-driven action potentials in CA3 pyramidal neurons from accelerating dye release. Presynaptic boutons were loaded by bath-applying 5 μM FM1-43 (Molecular Probes) in hypertonic ACSF supplemented with sucrose to 800 mOsm for 25 sec to selectively load the rapidly-recycling pool (RRP)[17,18], then returned to normal ACSF. Stimulus-induced destaining was measured after 30 min perfusion with dye-free ACSF, by bursts of 10 Hz bipolar stimuli (150 μs DC pulses) for 2 sec applied once each 30 sec. We fitted a single exponential to the first 6 fluorescence time course values, and decay time constants between groups compared by two-tailed Student’s t-test, as we have shown previously that the early release reflects vesicular release from the RRP prior to recycling and reuse of vesicles[17,18].

### Elevated Plus Maze

The elevated plus maze (EPM) is used as a general indicator of anxiety. The apparatus consists of two open arms (50 × 10 cm) and two closed arms (50 × 10 × 40 cm), connected by a center platform (10 × 10 cm) made of opaque dark grey plexiglass (Stoelting Inc., USA). The arms of the EPM are elevated 50 cm above the floor. Animals (n = 12-17) were placed in the center platform of the EPM, facing an open arm and allowed to explore the maze for 5 minutes. The middle point of the animal was used as the reference point to determine the position of a rat and recorded using behavioral software AnyMaze, Stoelting Co., Inc.). Percent of time spent in the open arms was calculated, where a decrease in percent of time spent in the open arms indicates an anxiety phenotype.

### Active Place Avoidance Learning Task

The active place avoidance task used was described previously by Burghardt et al.^19^. In this paradigm, mice are placed on a circular rotating platform that continuously turns clockwise at a speed of 1 rpm. Over several days and multiple trials (Fig. 3a), mice (n = 12-17) learn to identify the 60 shock zone guided by spatial markers on the walls surrounding the apparatus. Entrance into the shock zone triggered a brief constant foot-shock (500ms, 60Hz, 0.2mA) with an intershock interval of 1.5s that would cease upon leaving the shock zone. The middle point of the animal was used as the reference point to determine the position of a rat and recorded using behavioral software AnyMaze, Stoelting Co., Inc.). Using the same software, the number of shock-zone entries was measured, where a decrease in shock-zone entries indicates learning. During the initial pretraining trial (10 mins) when the shock was turned off, mice were allowed to habituate to the apparatus and showed no preference for any area of the platform. Subsequently, the shock was turned on and the mice completed 3 training sessions (10 mins each) per day for 3 days, followed by an extinction trial (10 mins) on the next day when the shock was turned off and the animals were allowed to ambulate freely into the zone previously associated with the shock. After extinction, a conflict variant task was performed in order to test cognitive flexibility. The shock zone was moved 180° from where the original shock zone was placed and 3 conflict-training sessions (10 mins each) were conducted per day for 2 days. Mice had to avoid the new shock zone which requires cognitive flexibility and is represented in this task by the simultaneous suppression of the learned condition response of avoiding the original shock zone and learning the new association of a foot-shock with a different zone.

**Figure 3.**
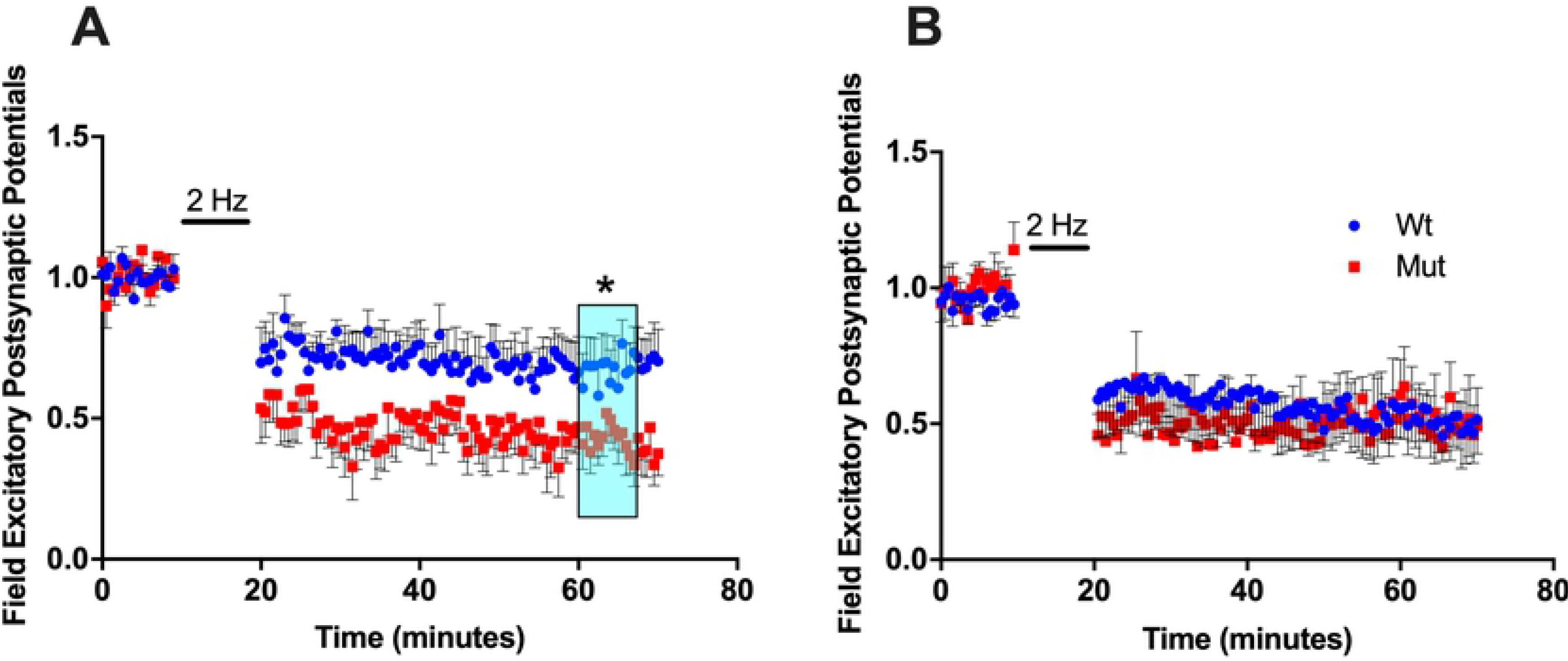
Changes in LTD recorded at Schaffer collateral-CA1 synapses in hippocampal slices from 1 and 4 month old SNAP-25 mutant mice. **(A)** LTD at 1 month of age at Schaffer collateral-CA1 synapses induced by a 2Hz LFS. SNAP-25b deficient mice (red squares, n=8) exhibited significantly greater LTD compared to controls (blue circles, n=7). Bar indicates application of LTD stimulus (data points not shown). (p<0.01; Student’s t-test). Each point is normalized to the averaged baseline and is mean ± SEM of n slices. **(B)** After prolonged absence of SNAP-25b, 4 month old SNAP-25b deficient mice (red squares, n=8) displayed LTD similar to wildtype littermates (blue circles, n=8). Bar indicates application of 2Hz/10mins stimulus (data points not shown). Each point is normalized to the averaged baseline and is mean ± SEM of n slices.

## Results

### Vesicular release probability is reduced in one month old SNAP-25b deficient mice

SNAP-25a and SNAP-25b are functionally different in their ability to facilitate exocytosis as they associate with the other core SNARE proteins in the SNARE complex[14]. Therefore, we analyzed whether SNAP-25a expressing mice exhibit functional differences in vesicular transmitter release properties at Schaffer collateral-CA1 synapses, as assessed by direct imaging of release of the styryl dye FM1-43 using two-photon laser scanning microscopy. By comparing gene targeted mice expressing only SNAP-25a to wildtype control mice, we compared the effects of a lack of SNAP-25b on release probability at glutamatergic Schaffer collateral terminals in hippocampal slices.

The technique of loading presynaptic vesicles with a membrane impermeable dye permits analysis of vesicle fusion dynamics in presynaptic terminals. FM1-43, preferentially loads into presynaptic vesicles. Once the dye is loaded into synaptic vesicles, it can only be released when vesicles fuse with the membrane and release their contents[17]. Two-photon analysis of FM1-43 makes it possible to directly image presynaptic rates of vesicular release[18]. To assess neurotransmitter release probability directly, Schaffer collateral presynaptic terminal vesicles were loaded with FM1-43 and the time course of fluorescent destaining in response to stimulus-evoked release of the dye was monitored using two-photon laser scanning microscopy[11,12]. Schaffer collateral axons were given a 2Hz stimulus train to evoke neurotransmitter release. Schaffer collateral terminals in field CA1 of hippocampal slices from SNAP-25a mice showed significantly slower neurotransmitter release kinetics compared to wildtype controls (Fig. 1a), as measured by a slower rate of fluorescence decay of FM1-43.

**Figure 1.**
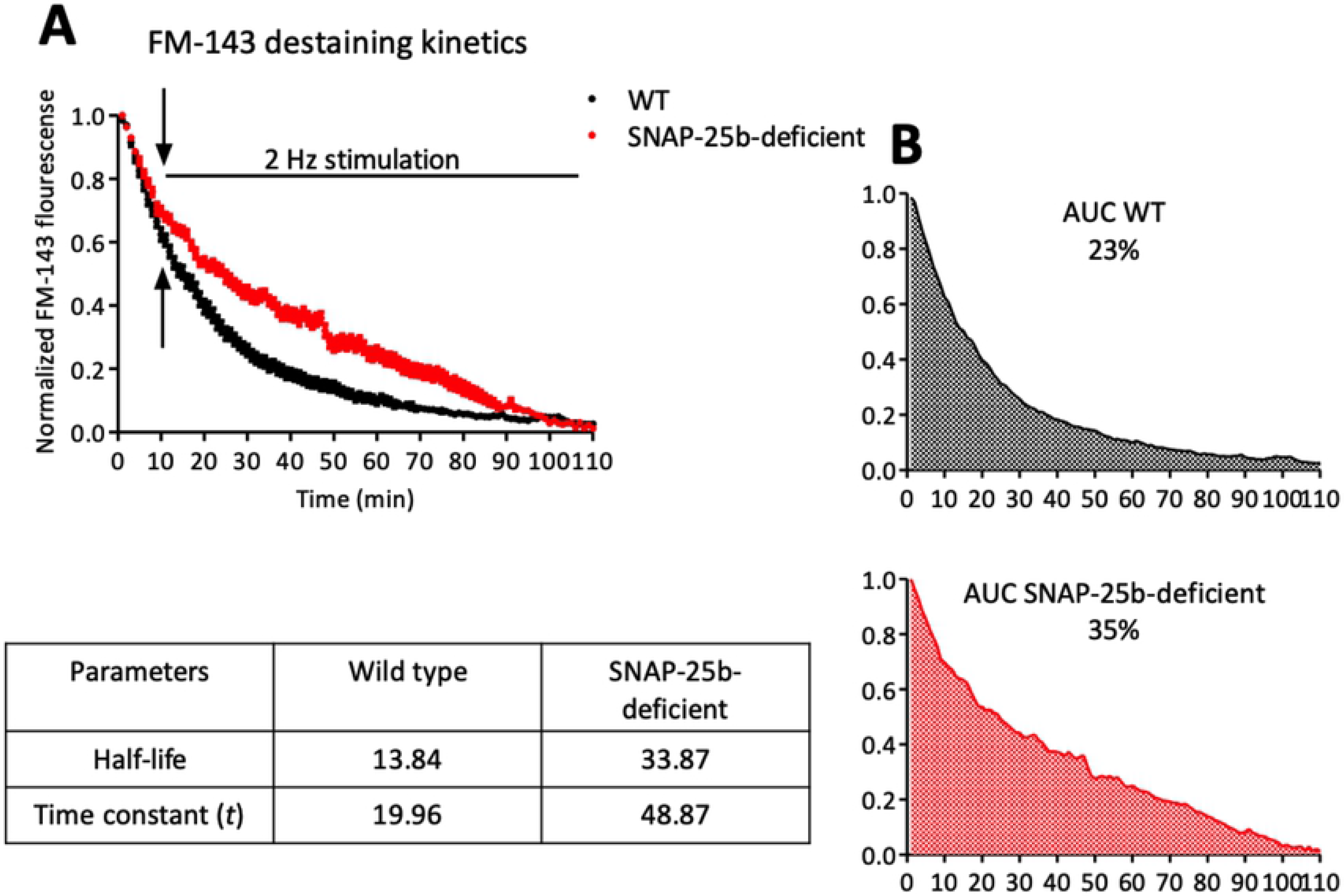
Presynaptic vesicular release from Schaffer collateral terminals in one-month old SNAP-25b deficient mice. **(A)** Fluorescence decay of vesicular dye FM1-43 from Schaffer collateral-CA1 presynaptic boutons following 2Hz stimulation (arrows represent the start of stimulation). SNAP-25b-deficient mice (red, n=14) showed a significantly slower rate of FM1-43 decay compared to wild type mice (black, n=20). Each point is a mean of n puncta. Fitting of the destaining curves with exponential one phase decay revealed significantly larger half-life and a larger decay time constant (*t*) values for SNAP-25b-deficient mice compared to the WT (p<0.05, Student’s t-test). (**B)** Calculating area under the curve (AUC) for the individual destaining curves also yielded higher AUC for the SNAP-25b-deficient mice (35%) compared to the WT (23%). Each point is a mean ± SEM of n slices.

### Long-term potentiation and long-term depression of synaptic strength

A high-frequency theta-burst stimulus (TBS) was used to induce stimulus-dependent LTP (sLTP) at Schaffer collateral-CA1 synapses in hippocampal slices from one and four month-old mice. At one month of age, sLTP was reduced in SNAP-25a mutant mice compared to age matched wildtype littermate controls (Fig. 2a). The reduced expression of LTP suggests an important role for the missing SNAP-25b isoform in the expression of LTP, though clearly there is LTP retained that does not require the presence of SNAP-25b.

**Figure 2.**
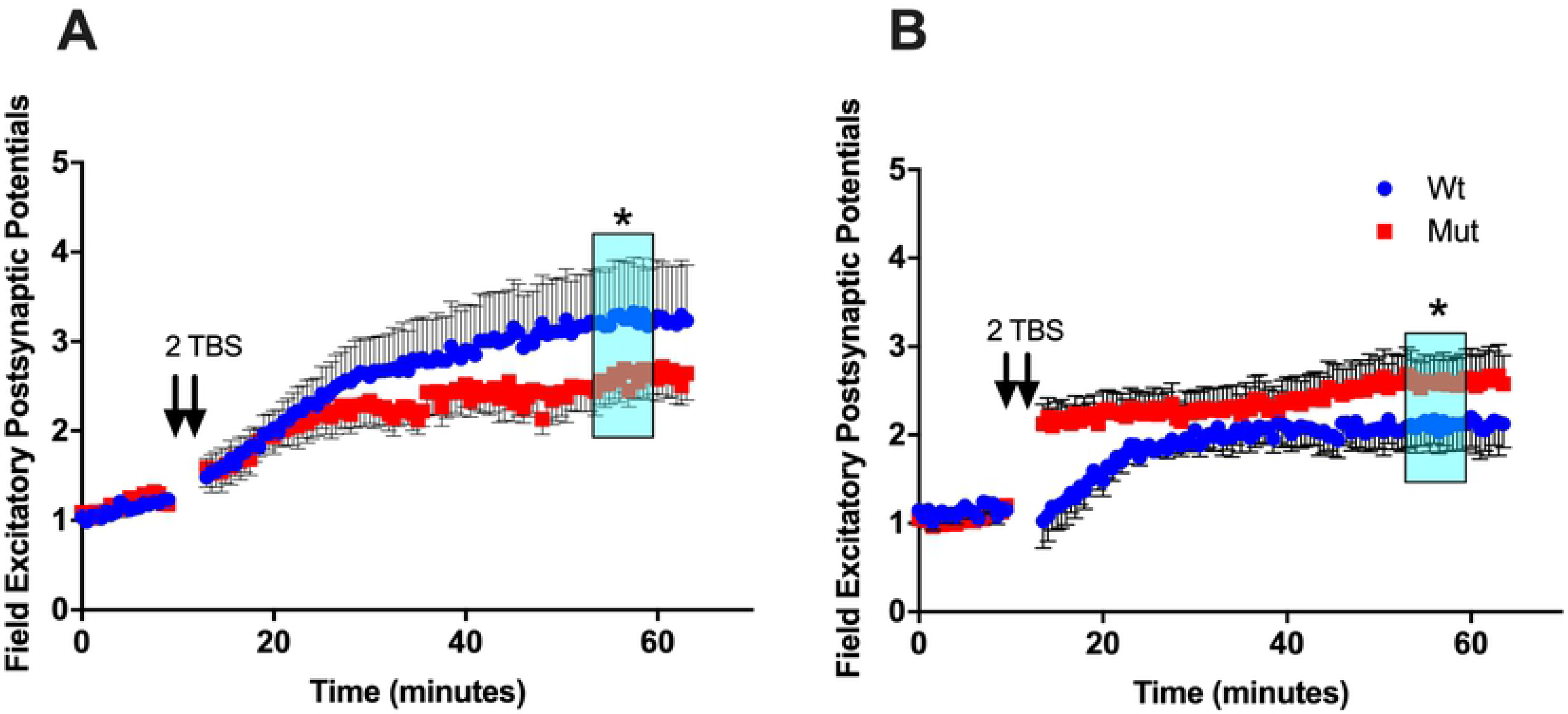
Alterations in stimulus-evoked sLTP detected in SNAP-25b deficient mice at 1 and 4 months of age. (**A)** Theta burst stimulation (TBS)-induced sLTP at 1 month in hippocampal Schaffer collateral synapses. Arrows indicate application of each TBS. Young SNAP-25b deficient mice expressed lower sLTP (red squares, n=13) compared to age-matched controls (blue circles, n=12). (p<0.01; Student’s t-test). Each point is normalized to the averaged baseline and is mean ± SEM of n slices. **(B)** Time course of LTP induced by TBS in littermate controls (blue circles, n=9) and SNAP-25b deficient mice (red squares, n=9) at slices from 4 month-old mice. Older SNAP-25b deficient mice show enhanced LTP with an altered PTP compared to littermate controls. (p<0.01; Student’s t-test). Each point is normalized to the averaged baseline and is mean ± SEM of n slices.

In contrast to observations in slices from one month old mice, four month old SNAP-25b deficient mice displayed elevated levels of LTP that were significantly larger than wildtype control mice (Fig. 2b). In addition to expressing augmented LTP, four month old SNAP-25b deficient mice exhibited larger rapid post-tetanic potentiation of fEPSPs (PTP) shortly after the HFS, in contrast with reduced PTP in one-month old mice. While one month old mice lacking SNAP-25b displayed reduced LTP the older animals, by four months of age, Schaffer collateral-CA1 synapses now exhibited enhanced LTP compare to age-matched controls, clear evidence for compensatory mechanisms at the presynaptic terminal that upregulate LTP in absence of the switch from SNAP-25a to SNAP-25b.

Due to the essential roles posited for both LTP and LTD in learning and memory, we also examined stimulus-evoked long-term depression of synaptic transmission (sLTD) in mice lacking SNAP-25b mice at both one and four months. At one-month of age, hippocampal slices from SNAP-25b deficient mice showed enhanced sLTD induced by a low-frequency stimulation (LFS; 2Hz/10min), compared to littermate controls (Fig. 3a). Taken together, our LTP and LTD data in 1-month-old animals suggests that SNAP-25a favors the expression of sLTD over sLTP in the developing brain, consistent with a previous study in which we showed that LTP at Schaffer collateral-CA1 synapses is reduced in both male and female SNAP-25b deficient mice at 1 month of age[15]. In contrast, four-month old SNAP-25b deficient mice displayed sLTD that did not differ from their littermate controls (Fig. 3b). These results also suggest compensatory effects can restore synaptic plasticity to control levels and balance in adult mice lacking SNAP-25b.

A recent study from our group[15] found, in an active avoidance spatial learning test, that this same one month old SNAP-25b deficient mouse shows impaired learning acquisition rates, and slower extinction of learning once acquired, suggesting weaker initial learning and greater flexibility for relearning new contingencies. Our current findings suggest that compensatory mechanisms have returned LTD and LTP to near control levels at 4 months of age in mice lacking the adult SNAP-25b isoform, leading to the question of whether the behavioral phenotypes we found in younger mice lacking SNAP-25b are also compensated in adulthood.

### Place avoidance spatial learning

To evaluate the behavioral phenotypes associated with a lack of SNAP-25b throughout the normal developmental period into adulthood, we used an active place avoidance assay developed by Fenton and colleagues, in which rodents are placed on a turning metal grid platform, and, using spatial cues, must learn to move to avoid a shock that is given when the animal enters one quadrant of the circular field. Littermate controls and SNAP-25b deficient mice were subject to multiday habituation, training trials, extinction and conflict discrimination described in Burghardt et al.[19]. In contrast to deficits in this learning, we have observed previously in one-month old mutant mice, adult mice lacking SNAP-25b showed no differences in the initial learning phase in days 1-3 (Fig. 4b). However, after extinction (day 4), when the shock zone was shifted 180° from its initial position, SNAP-25b deficient mice entered the new shock zone fewer times than controls, suggesting a more rapid relearning of the new shock zone location, a reflection of behavioral learning flexibility. No difference in anxiety-like behavior was detected in four-month old SNAP-25b deficient mice (Fig. 4c), using an elevated plus maze. Increased anxiety observed at one month in our previous study[15] was no longer detected in older SNAP-25b deficient mice. Finally, motor function as assessed by total path length was not different in SNAP-25b deficient mice and wildtype littermates (Fig. 4d). These data indicate that adult SNAP-25b-deficient mice have compensated for the impairments in synaptic plasticity and learning at 1 month of age, consistent with the shift from a reduction in LTP at on month of age, to larger LTP at 4 months of age, and the renormalization of the magnitude of LTD reached by adulthood.

**Figure 4.**
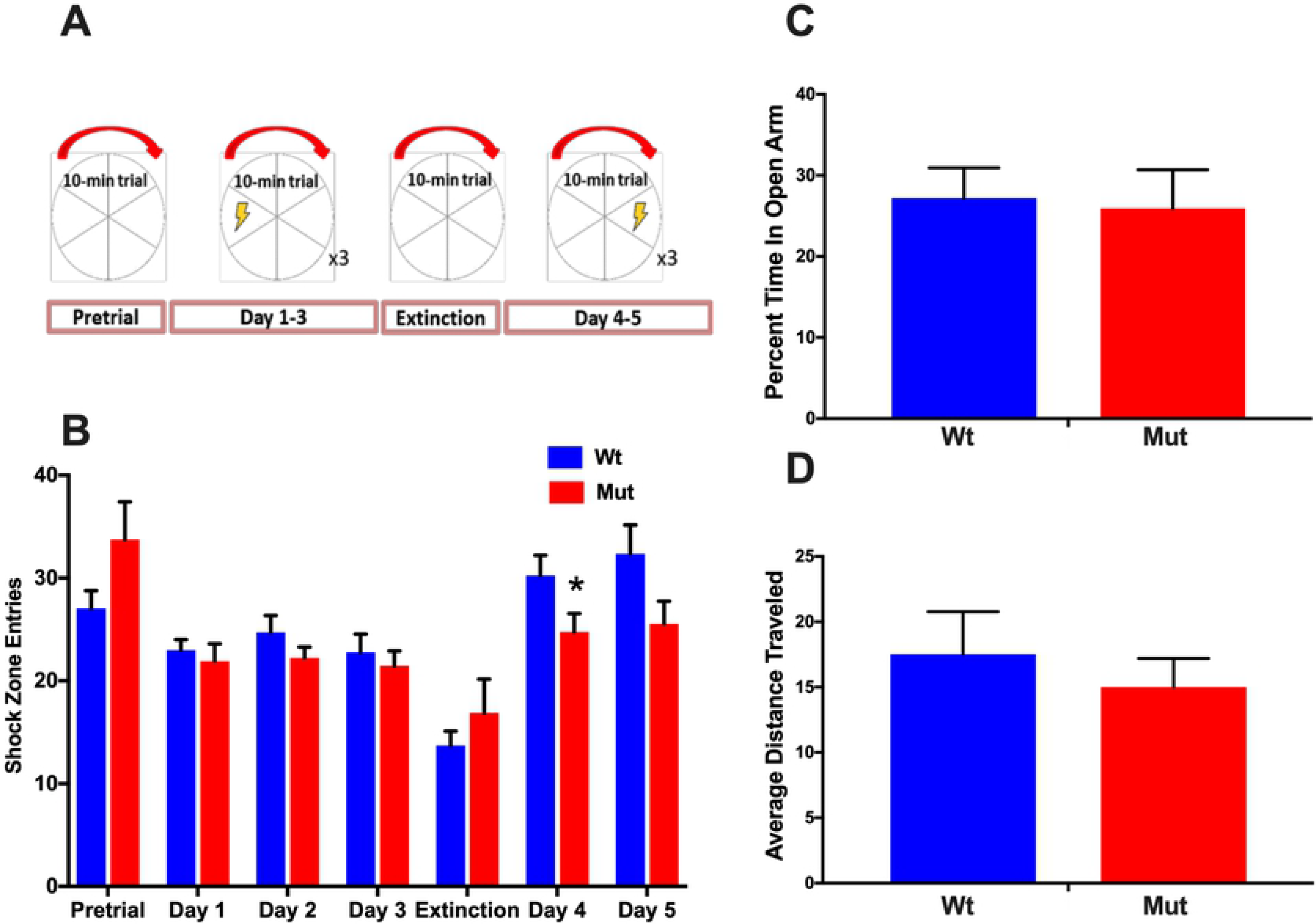
Active avoidance initially shows altered cognitive flexibility in adult SNAP-25b deficient mice with no difference in anxiety-like behaviors or locomotion. **A:** Active avoidance training schedule. Pretrial and Extinction: 1 - 10 min trial without shock. Initial learning: 3 days of 3 – 10 min trails/day with a left shock zone. Conflict learning: 2 days of 3-10 min trials/day with a 180° shifted shock region. **B:** Active avoidance assay average daily counts of shock zone entries. 4 month old SNAP-25b deficient mice (red bars, n=12) learned to avoid the original shock zone as well as controls (blue bars, n=17). After extinction, four-month old SNAP-25a mice initially learned faster and entered the new shock zone fewer times than wildtype littermates (day 4; p<0.05; Student’s t-test). **C:** 4 month old mice deficient in SNAP-25b showed no difference in time spent in the open arm of the EPM relative to controls. Therefore, no difference in anxiety-like behavior was detected. **D:** Older SNAP-25b deficient mice, compared to wildtype littermates, demonstrated no difference in general locomotion, indicated by total distance traveled. Each point is mean + SEM of n animals.

### Effects of learning acquisition on long-term synaptic plasticity

In a cohort of mice that went through active place avoidance behavioral testing, sLTP and sLTD were assessed post-training, to determine the impact of hippocampal-dependent training of place avoidance on hippocampal synaptic transmission and long-term activity-dependent plasticity in SNAP-25b deficient mice. Once mice completed all five days of the training paradigm, they were sacrificed and used for slice electrophysiology recordings. In contrast to untrained mice, one month old SNAP-25b deficient mice exhibited *enhanced* sLTP compared to littermate controls (Fig. 5a). Prior to training, the same SNAP-25b deficient mouse line had displayed reduced LTP at Schaffer collateral-CA1 synapses. While the response to HFS post-training was larger in SNAP-25b deficient mice at one month of age, the magnitude of sLTP was not altered by behavioral training in 4 month old SNAP-25b deficient mice (Fig. 5b).

**Figure 5.**
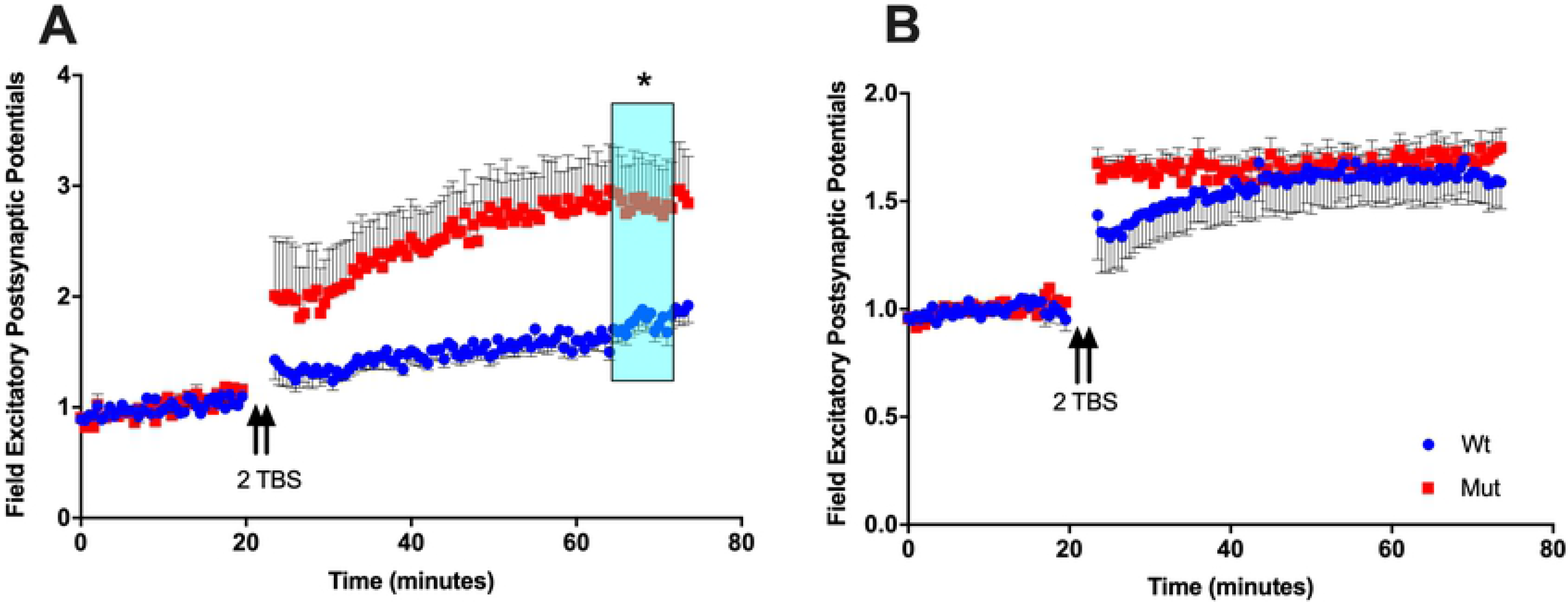
Hippocampal dependent training alters LTP in 1 month old SNAP-25b deficient mice. **A:** Time course of TBS (arrows; burst data points not shown) used to induce LTP in 1 month old mice. After completing an active avoidance assay, younger mice lacking SNAP-25b showed enhanced LTP (red squares, n=7) relative to controls (blue circles, n=11). p<0.01; Student’s t-test. Each point is normalized to the averaged baseline and is mean ± SEM of n slices. **B:** TBS LTP (arrows; burst data points not shown) recorded from CA1 stratum orients in 4 month old mice that had completed a multiday active avoidance training. SNAP-25b deficient mice (red squares, n=18) expressed LTP at levels similar to controls (blue circles, n=18). However, SNAP-25b deficient mice did show significantly larger PTP. Each point is normalized to the averaged baseline and is mean ± SEM of n slices.

Unlike adolescent mice lacking SNAP-25b, sLTD was altered in four-month old adult SNAP-25b deficient mice compared to wildtype littermate controls only after passive avoidance spatial learning task (Fig. 6b). Prior to training, four-month-old SNAP-25b deficient mice did not show altered sLTD, while younger mice showed the same enhancement in sLTD relative to controls both before and after hippocampal training. These data show that, in one month old mice lacking SNAP-25b, reductions in LTP and increases in LTD are both present, while by four months of age, compensatory mechanisms have returned LTD levels to normal. Nevertheless, the effect of learning acquisition on adult mice expressing only the immature SNAP-25a isoform was to upregulate both LTP and LTD at Schaffer collateral-CA1 synapses.

**Figure 6.**
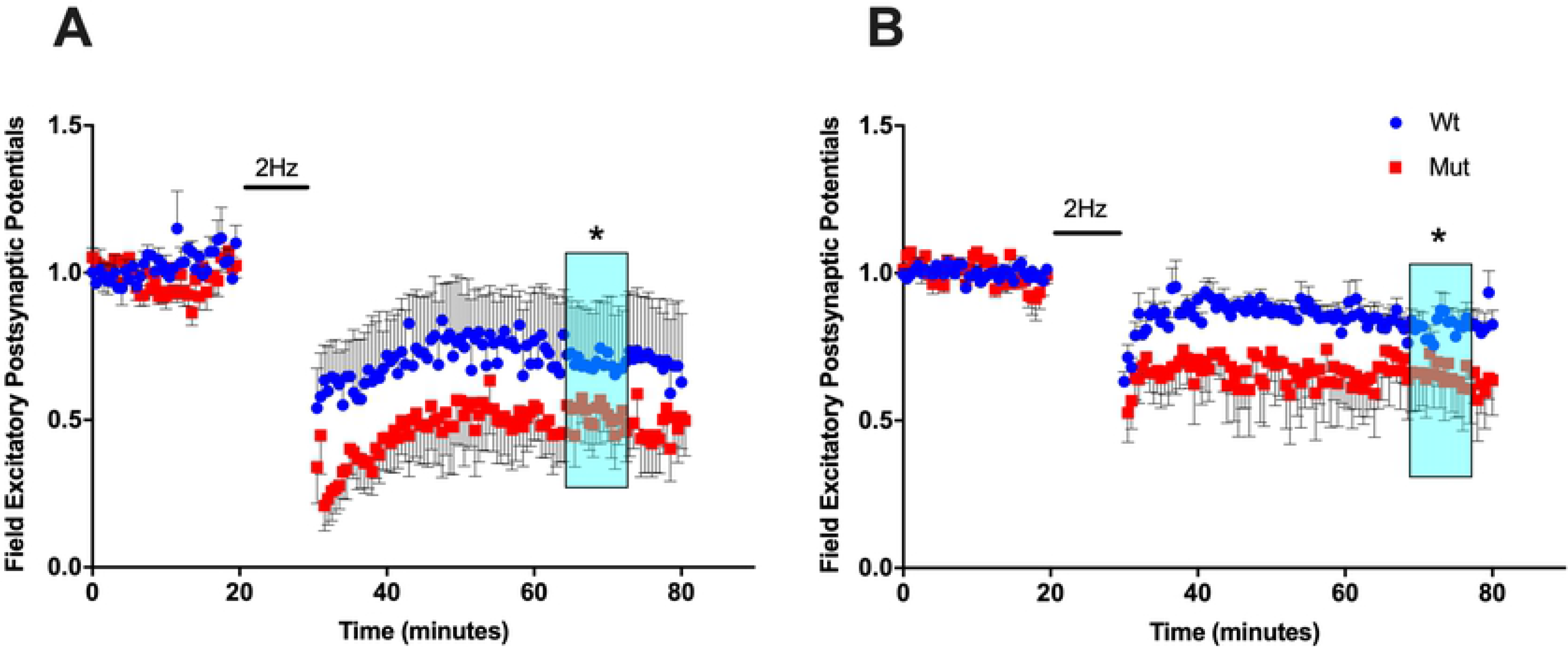
LTD is enhanced in young and old SNAP-25b deficient mice post hippocampal dependent training. **(A)** 1 month old SNAP-25b deficient mice (red squares, n=7) exhibited significantly larger sLTD compared to wildtype littermates (blue circles, n=4). (p<0.01; Student’s t-test). Bar indicates 2Hz/10mins LTD stimulus application. Each point is normalized to the averaged baseline and is mean ± SEM of n slices. (**B)** 4 month old SNAP-25b deficient mice (red squares, n=6) also exhibited greater sLTD than wildtype controls (blue circles, n=13, p<0.05; Student’s t-test). Bar indicates 2Hz/10min LTD LFS stimulus train. Each point is normalized to the averaged baseline and is mean ± SEM of n slices.

### mGluRII and NMDA receptor-dependent LTD of synaptic transmission

Given the role of SNAP-25 in regulating neurotransmitter release, we hypothesized that changes in activation of glutamate receptors promote presynaptic long-term activity-dependent plasticity might be responsible for differences in synaptic plasticity resulting from a lack of SNAP-25b. Multiple reports indicating an essential role for SNAP-25 in transient presynaptic suppression of transmitter release by the 5-HT serotonin receptor at lamprey synapses[23,24], and in mGluRII-dependent chemical and stimulus-evoked LTD at Schaffer collateral-CA1 hippocampal synapses^25^. Evidence points to an interaction between the C-terminus of SNAP-25 with the G-protein Gβγ being necessary for suppression of neurotransmitter release at all of these synapses[24–27]. C-terminus cleavage of SNAP-25 by BoNT/A prevents Gβγ and SNAP-25 interaction necessary for depression of transmitter release, as does infusion of a c-terminal fragment of SNAP-25 that binds to Gβγ, in both lamprey reticulospinal synapses and rat Schaffer collateral synapses[23,25]. Given the role of these glutamate receptors in the induction of LTD of presynaptic transmitter release, we tested whether there are differences in the expression of mGluRII or NMDAR cLTD produced by a lack of SNAP-25b. To evaluate the involvement of each form of presynaptic cLTD and how they interface with SNAP-25 to affect presynaptic vesicular release, mGluRII- and NMDAR-dependent LTD were selectively induced in hippocampal slices from one and four-month old mice lacking SNAP-25b, by bath application of either the mGluRII agonist DCG-IV (25μM) or NMDA (20μM).

In mGluRII cLTD, lack of SNAP-25b did not alter the amplitude of mGluRII cLTD in either one-(Fig. 7a) or four-month old (Fig. 7b) SNAP-25b deficient mice compared to wildtype littermate controls, indicating that molecular mechanisms downstream of synaptic stimulation that underlie the induction of mGluRII-dependent presynaptic LTD of transmitter release are not differentially regulated by the two isoforms of SNAP-25. NMDAR cLTD also exhibited the same amplitude of fEPSP in one-month old SNAP-25a mutant mice as in control mice (Fig. 8). If components of the NMDAR cascade interact with SNAP-25, this interaction does not appear to be influenced by a lack of SNAP-25b in adolescent mice. Further, we found that NMDAR-dependent LTD could not be elicited in either mutant or control mice by bath application of NMDA at four months of age, indicating that synaptic stimulation is an essential component in the induction of LTD in adult mice, since a significant component of sLTD is blocked by NMDAR antagonists. Taken together, these findings suggest that the a and b isoforms of SNAP-25 differentially regulate the induction of LTD through regulation of the magnitude and patterns of synaptic stimulation and its effects on presynaptic terminals that require synaptic activity, rather than by altering the downstream activation of glutamate receptors either postsynaptically or presynaptically.

**Figure 7.**
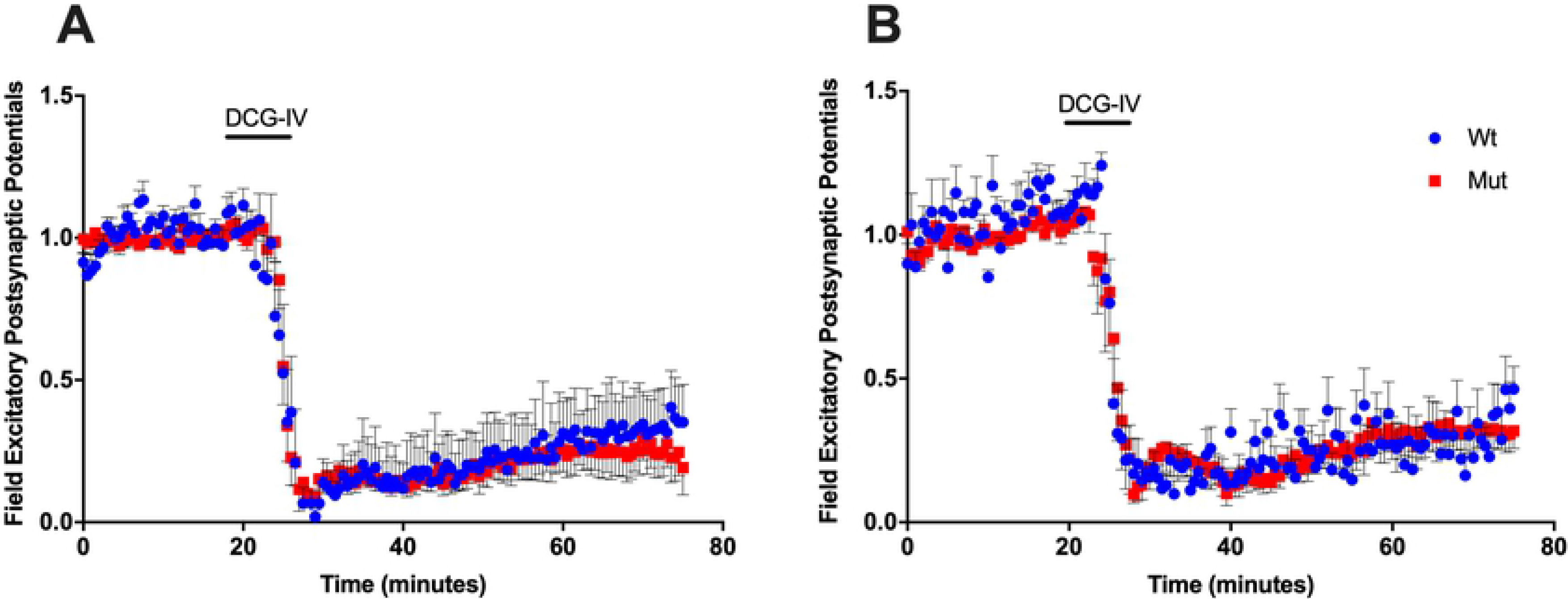
1 and 4 month old SNAP-25 mutant mice show no differences in mGluRII dependent cLTD. Time course of mGluRII cLTD induced by the receptor agonist, DCG-IV (25μM; solid bar = 5 mins at 2mL/min) at Schaffer collateral-CA1 synapses. Each point is normalized to the averaged baseline and is mean ± SEM of n slices. **A:** In 1 month old SNAP-25b deficient hippocampal slices (red squares, n=11), mGluRII LTD was not altered compared to littermate controls (blue circles, n=7) **B:** In 4 month old SNAP-25b deficient (red squares, n=7) and wildtype (blue circles, n=9) mice, mGluRII LTD did not differ in magnitude.

**Figure 8.**
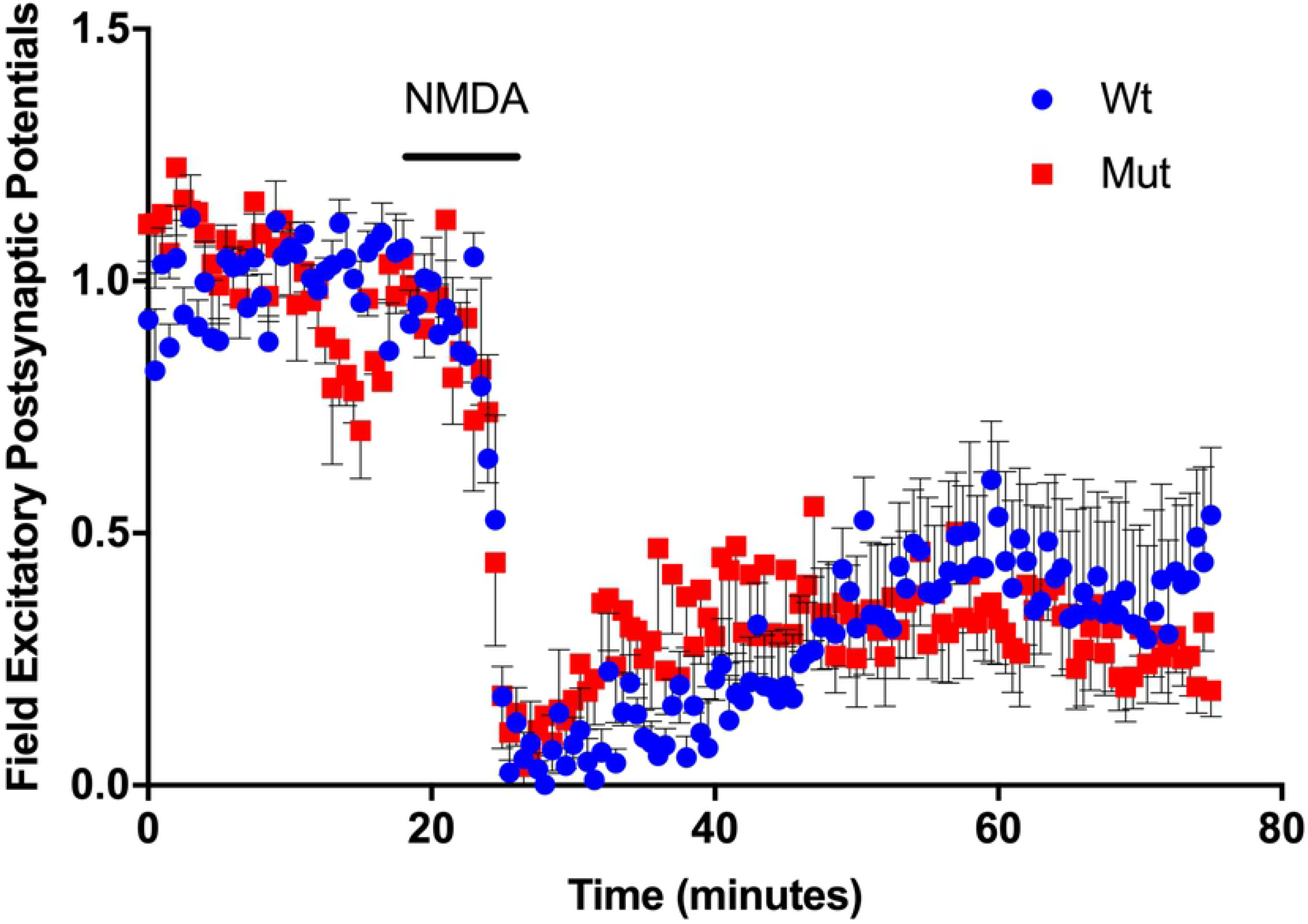
Differences observed in synaptic plasticity in one month old SNAP-25b deficient mice were not due to alterations in NMDAR response to glutamate. The presence of SNAP-25a in mutant mice (red squares, n=5) did not alter the extracellular amplitude of NMDA-induced cLTD (20μM; solid bar = 5 mins at 2mL/min) compared to wildtype littermates (blue circles, n=7) at Schaffer collateral-CA1 synapses. Each point is normalized to the averaged baseline and is mean ± SEM of n slices.

## Discussion

SNAP-25 is an essential SNARE complex protein and a regulatory target for the expression of short and long-term plasticity of presynaptic transmitter release. SNAP-25 exists in alternative splice isoforms SNAP-25a and b, with the a isoform predominating in immature rain, while the b isoform is the major one in the adult brain. The individual contributions of each isoform to the regulation of synaptic transmission and long-term synaptic plasticity is still unclear. Previous work suggests that there may be a temporal association with the expression of SNAP-25 isoforms and the type of activity-dependent synaptic plasticity that can be elicited. Given the importance of LTP and LTD in activity-dependent networks that underlie learning and memory, we assessed alterations in LTP, LTD and behavioral learning at Schaffer collateral-CA1 synapses in hippocampal slices in a gene targeted mouse that lacks SNAP-25b, compared to wildtype mice that exhibit normal levels and timing of SNAP-25a and b expression.

The reduced level of LTP and enhanced magnitude of LTD expressed in one-month old SNAP-25b deficient mice relative to littermate controls suggests that SNAP-25a favors the induction of LTD over LTP, and is consistent with our previous study showing that both male and female SNAP-25b deficient mice exhibit significantly smaller LTP[15]. By preventing a down regulation of LTD during development, the absence of SNAP-25b could prevent the shift in synaptic plasticity favoring larger LTP that normally occurs around 3 weeks of age[9]. During the first 8 weeks of development, wildtype mice display a dramatic increase in SNAP-25b mRNA expression, and this corresponds to a developmental window for complete maturation of synaptic connections^9^. Therefore, the loss of SNAP-25b could alter synaptic maturation by modifying activity-dependent synaptic plasticity. In this study, SNAP-25a mice showed a delayed progression in normal expression levels of LTP and LTD. It is probable that, without the dramatic increase in SNAP-25b which markedly decreases the ratio of SNAP-25a to SNAP-25b, one-month old SNAP-25b deficient mice were more like younger wildtype mice, where the ratio of SNAP-25a/SNAP25b is still high^2^, and LTD is favored over LTP, consistent with our observations in gene targeted mice expressing only SNAP-25a. This observation is in line with previous studies of SNAP-25a, reporting that the adolescent isoform is less efficient at coordinating vesicle release[9,20,28]. The results of our study support the conclusion that the switch from SNAP-25a to SNAP-25b may be a key mediator of the developmental switch in relative magnitudes of LTD and LTP seen in the hippocampus of developing mice. It would be useful to develop a complementary mouse that expresses only SNAP-25b throughout the lifetime of the mouse, to test the prediction that LTD would be impaired early in development, and LTP would emerge prematurely in ways that would alter normal brain development.

By four months of age, SNAP-25b deficient animals appeared to have over-compensated for the lack of SNAP-25b. In the process of up regulating the strength of LTP to match the levels of LTP in adult wildtype mice, hippocampal Schaffer collateral-CA1 synapses in SNAP-25b deficient mouse slices exhibited enhanced LTP relative to wildtype controls. It is intriguing that these synapses also showed larger PTP immediately after HFS. The increase in PTP may be due to increased residual calcium remaining in the presynaptic terminal after HFS, suggesting the possibility of alterations in presynaptic calcium regulation that increase vesicular release immediately after HFS. An enhancement of both PTP and LTP could also signify an up regulation of NMDAR-gated postsynaptic currents. In particular, NR2B subunit containing NMDARs can be up regulated, resulting in enhanced LTP[29,30].

In contrast, levels of LTD in these older SNAP-25b deficient mice were similar to controls, suggesting that, if NR2B NMDAR subtypes are lesser participants in the induction of LTD, presynaptic sites of regulation may account for compensatory shifts in expression of LTD in older mice. With regard to the amplitude of LTD, SNAP-25b lacking mutant mice were able to accurately compensate for the enhancement seen in younger mice, bringing LTD back to levels observed in littermate controls. Taken together, findings at one and four months of age confirm the hypothesis that the development shift from SNAP-25a to increased SNAP-25b is an important factor in regulating the relative magnitude of sLTP and sLTD in the developing and adult brain.

It is unclear whether compensatory mechanisms underlying shifts in LTP and LTD result from a lack of SNAP-25b, or over-expression of SNAP-25a. Although the adolescent a isoform is less effective at coordinating vesicle fusion, it can rescue SNAP-25-/- neurons in culture and coordinate synchronous stimulus evoked release[28]. The absence of SNAP-25b may promote enhanced functionality or increased recruitment of SNAP-25a or altered association of SNAP-25a with accessory proteins, as mice age. A previous study has shown that the major functional differences between SNAP-25a and SNAP-25b are due to two residues at the C-terminus, and that these residues alter protein-protein interactions[10]. Additionally, Munc 18-1, an accessory SNARE protein which promotes SNARE complex formation, also interacts with SNAP-25 and binds to SNAP-25b more easily than SNAP-25a[31]. Differential interaction of SNAP-25 isoforms with Munc-18-1 could potentially alter the rate of SNAP-25a incorporation into SNARE complexes during vesicle priming. As mice age and more SNAP-25a is incorporated into SNARE complexes, more vesicles may be accessible for synaptic transmission and account for the compensatory effects seen in older SNAP-25b deficient mice. To test this possibility, it would be helpful to investigate changes in the function and size of the RRP in SNAP-25b deficient mice at one and four months, using imaging of FM1-43 release and electron microscopy. Alternatively, these compensatory mechanisms may be postsynaptic in origin, involving changes such as up-regulation of NR2B-NMDAR expression[30] to compensate for continued impairments in presynaptic function. In future experiments, it will be essential to use whole-cell patch-clamp recording of postsynaptic NMDA currents to probe for changes in NMDAR currents as mice age.

To assess whether hippocampal dependent learning and memory is affected by lack of the adult SNAP-25b isoform throughout development, hippocampal-dependent spatial learning was evaluated in four month old SNAP-25b deficient mice using an active avoidance assay. Four month old SNAP-25b deficient mice initially showed no differences in acquisition of learned place avoidance, but they did show an enhanced conflict learning immediately after extinction, i.e., a more rapid shift away from a previous shock zone to learn a new zone. On the second day of conflict reversal learning, there was only a non-significant trend towards fewer shock zone entries for the mice lacking SNAP-25b. The possibility exists that, since older SNAP-25b deficient mice did not experience the initial delay in learning we observed in younger mice, their underlying memory deficit had largely disappeared as compensatory mechanisms were expressed during development. Given that, at one month of age, this same mouse line showed enhanced cognitive learning, we hypothesize that enhanced LTD in these adolescent animals accounted for their performance during the relearning phase of the test, and both LTD and learning had returned to normal in adulthood.

To address any underlying memory deficit not detected by the active avoidance assay, additional behavioral learning and memory paradigms will be needed to test hippocampus-dependent and independent forms of learning, retention, and relearning, to determine if behavioral flexibility is altered by the subtle differences in presynaptic function afforded by the isoforms of SNAP-25. For example, it is possible to increase the complexity of the spatial avoidance learning task by adding a rotating shock segment with the stationary segment during the initial learning phase[19].

Mice expressing only SNAP-25a exhibit a higher number of doublecortin positive precursor cells[11], which implies a higher rate of neurogenesis in these mice. Increased neurogenesis may lead to improved performance on cognitive flexibility tasks[19], and could serve as an alternative explanation for the results of observed here. Prior to training in the active avoidance task, mice were monitored for anxiety. The time spent in the open potion of the elevated plus maze relative to total time in the apparatus was used as an indicator of anxiety state. At four months of age, SNAP-25b deficient animals were no different from controls and showed comparable levels of locomotion, indicating that our spatial learning findings cannot be attributed to alterations in motivation.

Some of the same mice that went through active avoidance behavioral testing, referred to as trained mice, were then monitored for changes in sLTP and sLTD. Trained one month old SNAP-25b deficient mice expressed enhanced, instead of reduced, LTP, but still showed the larger LTD relative to littermate controls that we observed in untrained mice. The relative switch in the strength of LTP in trained versus untrained mice lacking SNAP-25b suggests that the act of training functionally improved synaptic plasticity, or that SNAP-25b deficient mice demonstrate less LTP elicited by active avoidance training itself, leaving more room below ceiling levels for potentiation to be elicited by HFS.

In contrast to adolescent mice, adult trained mice exhibited LTP similar in magnitude to controls, but LTD was still significantly enhanced in magnitude. In untrained SNAP-25b deficient mice at one month of age, LTP was lower than controls. However, by four months of age untrained SNAP-25b deficient mice show enhanced LTP. Since adolescent untrained SNAP-25b deficient mice exhibited smaller LTP than controls, compensatory mechanisms may have been activated that resulted in overshooting the normal homeostatic set point in naïve mice, leading to larger LTP in untrained SNAP-25b deficient mice. Conceivably, the training paradigm could have activated cascades that reset synapses to normal levels of LTP, and a consequence of this compensation to adjust LTP may have been responsible for the enhancement of LTD. Moreover, the difference seen in SNAP-25b deficient mice post training could have been due to a shift in the ability to regulate the sliding threshold of the synapse. If the sliding threshold scale is being altered in SNAP-25b deficient mice as a compensatory mechanism to accommodate the deficits in synaptic plasticity, it could imply that SNAP-25 function contributes to a synapses’ ability to regulate metaplasticity or the threshold for changing synaptic strength. In either case, it would be interesting to further investigate the compensatory mechanisms that may account for these observations, including whole-cell patched clamp analysis of postsynaptic NMDAR currents, functional analysis of presynaptic release probability, vesicular release rates of FM1-43, and electron microscopy of Schaffer collateral presynaptic terminals to quantify the relative number of docked vesicles.

To fully evaluate the effects of a lack of SNAP-25b on baseline transmitter release, we measured presynaptic neurotransmitter release with FM1-43. The FM1-43 data described in this study indicates a slower rate of neurotransmitter release associated with multiple forms of stimulus and glutamate receptor-evoked LTD. The decreased rate of release seen in mice lacking SNAP-25b implies a lower release probability following the induction of LTD. PPF is also an indicator of presynaptic neurotransmitter release mechanisms. Larger PPF means that a smaller number of vesicles released their content after the first pulse due to a lower release probability, resulting in larger PPF in response to a second stimulus[17,32]. In a recent study[15], we found that PPF in SNAP-25b deficient mice was significantly larger than in SNAP-25b expressing wildtype mice, also suggesting that release probability is lower in mice expressing only SNAP-25a. In this earlier study[15], we also found that I/O ratios were unchanged, suggesting that SNAP-25b deficient mice had homeostatically compensated for lower release probabilities by upregulating postsynaptic sensitivity to glutamate.

The results described above regarding baseline synaptic transmission may put behavioral findings associated with SNAP-25a into context. It is consistent with previous studies[20,28] showing that SNAP-25a containing SNARE complexes are slower to release FM1-43, indicating lower release probability. Impaired transmitter release mechanisms may result in memories that are not as persistent and stable as wildtypes, making them more susceptible to reversal in conflict learning assays. Reduced transmitter release probability is suggested to impair the induction of LTP, resulting in weaker synaptic strengthening and memory formation. Therefore, extinction in adult SNAP-25b deficient mice may proceed faster than in control mice that express adult SNAP-25b. It is noteworthy that the I/O relationship is similar to controls[15], suggesting that compensatory postsynaptic changes may have been activated in the face of basal reductions in release probability.

The absence of SNAP-25b did not alter the amplitude of mGluRII-dependent LTD measured at Schaffer collateral-CA1 synapses when DCG-IV was bath applied, or NMDAR-dependent LTD when NMDA was bath applied, to hippocampal slices from these mice, compared to littermate controls at four months of age. This suggests that SNAP-25a and Gβγ can interact to induce cLTD in adult SNAP-25b-deficient mice with similar affinity to wildtype mice during both adolescence and adulthood. These results are not consistent with recent immunoprecipitation data showing that SNAP-25a-containing SNARE complexes associate with Gβγ 50% less than SNAP-25b-containing complexes[25]. Since, even if Gβγ is less likely to interact with SNAP-25a, a difference in chemically-induced NMDA or mGLuRII-dependent LTD was not detected, it is possible that lower levels Gβγ and SNAP-25 interaction may still be sufficient to permit expression of normal magnitude NMDAR and mGluRII cLTD. These results also imply that the altered synaptic plasticity induced by stimulus in young and old mutant mice (SNAP-25a) versus wild type (normal SNAP-25b) mice may have been in response to altered presynaptic activity, since we observed no difference in fEPSPs when mGluRII-LTD was expressed in SNAP-25b deficient mice. Our previous work in rats has shown that presynaptic infusion of Gβγ scavenging peptides, ct-SNAP-25 and structurally distinct mSIRK, can each occlude mGluRII LTD[31]. Therefore, Gβγ binding to SNAP-25a in mice expressing only this isoform may show larger sLTD compared to littermate controls, given evidence that SNAP-25a favors LTD. An important next study will be to evaluate the vesicular release of FM1-43 from SNAP-25a mutants after expressing each form of chemically-induced LTD.

In summary, one role of the switch from expression of the immature SNAP-25a isoform to the adult SNAP-25b isoform seems to be to down-regulate the predominance of LTD of synaptic strength in early activity-dependent long-term plasticity, favoring an up-regulation of LTP that promotes superior learning acquisition and retention, but reducing behavioral flexibility when learning contingencies change.

## Acknowledgements

This work was supported by grants from the Sven Mattssons Foundation (CB), The Swedish Brain Foundation (CB), and NIH NRSA Fellowship (KRG).

## References

1. Söllner, T., Bennett, M. K., Whiteheart, S. W., Scheller, R. H. & Rothman, J. E. A protein assembly-disassembly pathway in vitro that may correspond to sequential steps of synaptic vesicle docking, activation, and fusion. Cell 75, 409–418 (1993).

2. Jahn, R. & Scheller, R. H. SNAREs--engines for membrane fusion. Nat Rev Mol Cell Biol 7, 631–643 (2006).

3. Südhof, T. The molecular machinery of neurotransmitter release (Nobel Lecture). Angew Chem Int Ed Engl 53, 12696–12717 (2014).

4. Fang, Q. et al. The role of the C terminus of the SNARE protein SNAP-25 in fusion pore opening and a model for fusion pore mechanics. Proc Natl Acad Sci U S A 105, 15388–15392 (2008).

5. Osen-Sand, A. et al. Inhibition of axonal growth by SNAP-25 antisense oligonucleotides in vitro and in vivo. Nature 364, 445–448 (1993).

6. Pan, Q., Shai, O., Lee, L. J., Frey, B. J. & Blencowe, B. J. Deep surveying of alternative splicing complexity in the human transcriptome by high-throughput sequencing. Nat Genetics 40, 1413–1415 (2008).

7. Wang, E.T. et al. Alternative isoform regulation in human tissue transcriptomes. Nature 456,470–476 (2008).

8. Grosse, G. et al. SNAP-25 requirement for dendritic growth of hippocampal neurons. J Neurosci Res 56, 539–546 (1999).

9. Bark, I.C., Hahn, K.M., Ryabinin, A.E. & Wilson, M.C. Differential expression of SNAP-25 protein isoforms during divergent vesicle fusion events of neural development. Proc Natl Acad Sci U S A 92, 1510–1514 (1995).

10. Nagy, G. et al. Alternative splicing of SNAP-25 regulates secretion through nonconservative substitutions in the SNARE domain. Molecular biology of the cell 16, 5675–5685 (2005).

11. Johansson, J.U. et al. An ancient duplication of exon 5 in the Snap25 gene is required for complex neuronal development/function. PLoS Genetics 4(11), e1000278 (2008).

12. Valladolid-Acebes, I. et al. Replacing SNAP-25b with SNAP-25a expresion results in metabolic disease. Proc Natl Acad Sci (USA) 112,4326–4335 (2015).

13. Bolshakov Y.Y. & Siegelbaum, S.A. Postsynaptic induction and presynaptic expression of hippocampal long-term depression. Science 264,1148–1152 (1994).

14. Dudek, S.M. & Bear, M.F. Bidirectional long-term modification of synaptic effectiveness in the adult and immature hippocampus. J Neurosci 13,2910–2918 (1993).

15. Irfan, M. et al. SNAP-25 isoforms differently regulate synaptic transmission and long-term synaptic plasticity at central synapses. Scientific Reports, in review (2019).

16. Bark, C. et al. Developmentally regulated switch in alternatively spliced SNAP-25 isoforms alters facilitation of synaptic transmission. J Neurosci 24, 8796–8805 (2004).

17. Stanton, P.K. et al. Long-term depression of presynaptic release from the readily-releasable vesicle pool induced by NMDA receptor-dependent retrograde nitric oxide. J Neurosci 23,5936–5944 (2003).

18. Stanton, P.K. et al. Imaging LTP of presynaptic release of FM1-43 from the rapidly-recycling vesicle pool at Schaffer collateral-CA1 synapses in rat hippocampal slices. Eur J Neurosci 22,2451–2461 (2005).

19. Burghardt, N.S., Park, E.H., Hen, R. & Fenton, A.A. Adult-born hippocampal neurons promote cognitive flexibility in mice. Hippocampus 22, 1795–1808 (2012).

20. Sørensen, J.B. et al. Differential control of the releasable vesicle pools by SNAP-25 splice variants and SNAP-23. Cell 114, 75–86 (2003).

21. Wu, L.G. & Betz, W.J. Nerve activity but not intracellular calcium determines the time course of endocytosis at the frog neuromuscular junction. Neuron 17, 769–779 (1996).

22. Winterer, J., Stanton, P.K. & Müller, W. Direct monitoring of vesicular release and uptake in brain slices by multiphoton excitation of the styryl FM 1-43. Biotechniques 40, 343–351 (2006).

23. Blackmer, T. et al. G protein betagamma subunit-mediated presynaptic inhibition: regulation of exocytotic fusion downstream of Ca2+ entry. Science 292, 293–297 (2001).

24. Blackmer, T. et al. G protein betagamma directly regulates SNARE protein fusion machinery for secretory granule exocytosis. Nat Neurosci 8, 421–425 (2005).

25. Zhang, X.L., Upreti, C. & Stanton, P.K. Gβγ and the C terminus of SNAP-25 are necessary for long-term depression of transmitter release. PLoS One 6, e20500 (2011).

26. Gerachshenko, T. et al. Gbetagamma acts at the C terminus of SNAP-25 to mediate presynaptic inhibition. Nature neuroscience 8, 597–605 (2005).

27. Photowala, H., Blackmer, T., Schwartz, E., Hamm, H.E. & Alford, S. G protein betagamma-subunits activated by serotonin mediate presynaptic inhibition by regulating vesicle fusion properties. Proc Natl Acad Sci U S A 103, 4281–4286 (2006).

28. Delgado-Martínez, I., Nehring, R.B. & Sørensen, J.B. Differential abilities of SNAP-25 homologs to support neuronal function. J Neurosci 27, 9380–9391 (2007).

29. Chen, Y. et al. Hippocampal NR2B-containing NMDA receptors enhance long-term potentiation in rats with chronic visceral pain. Brain Res 1570, 43–53 (2014).

30. Müller, L., Tokay, T., Porath, K., Köhling, R. & Kirschstein, T. Enhanced NMDA receptor-dependent LTP in the epileptic CA1 area via upregulation of NR2B. Neurobiol Dis 54, 183–193 (2013).

31. Daraio, T., Valladolid-Acebes, I., Brismar, K. & Bark, C. SNAP-25a and SNAP-25b differently mediate interactions with Munc18-1 and Gβγ subunits. Neurosci Lett 674, 75–80 (2018).

32. Thomson, A.M. Facilitation, augmentation and potentiation at central synapses. Trends Neurosci 23, 305–312 (2000).

